# PlantC2U: Deep learning of cross-species sequence landscapes predicts plastid C-to-U RNA editing in plants

**DOI:** 10.1101/2023.05.18.541274

**Authors:** Chaoqun Xu, Jing Li, Ling-Yu Song, Ze-Jun Guo, Shi-Wei Song, Lu-Dan Zhang, Hai-Lei Zheng

## Abstract

In plants, C-to-U RNA editing is mainly occurred in the plastids and mitochondria transcripts, which contributes to complex transcriptional regulatory network. More evidences reveal that RNA editing plays critical roles in plant growth and development. However, RNA editing sites accurately detected by transcriptome sequencing data alone are still challenging. In the present study, we developed PlantC2U, which is a convolutional neural network to predict plastid C-to-U RNA editing based on the genomic sequence. PlantC2U achieves over 95% sensitivity and 99% specificity, which outperforms random forest and support vector machine. PlantC2U not only further checks RNA editing sites from transcriptome data to reduce the possible false positives, but also assesses the effect of different mutations on C-to-U RNA editing status based on the flanking sequences. Moreover, we found the patterns of tissue-specific RNA editing in mangrove plant *Kandelia obovata*, and observed reduced C-to-U RNA editing rates in cold stress response of *K. obovata*, suggesting their potential regulatory roles in the plants stress adaption. In addition, we present RNAeditDB, available online at https://jasonxu.shinyapps.io/RNAeditDB/. Together, PlantC2U and RNAeditDB would help researchers explore the RNA editing events in plants and thus would be of broad utility for the plant research community.

**Highlight:** We develop a convolutional neural network based deep learning, PlantC2U program, which help researchers explore the plastids C-to-U RNA editing events in plants and thus would be of broad utility for the plant research community.

## Introduction

RNA editing is a universal phenomenon that can modify the RNA sequence compositions through nucleotides deletion, insertion and other base substitution, which is an important post-transcriptional process (Brennicke *et al*., 1999). In plants, the most major type of RNA editing in the organellar mRNA is cytidine to uridine (C-to-U) transition (Small *et al*., 2020), which was first reported in mitochondria in 1989 (Hiesel *et al*., 1989), and chloroplast in 1991 (Hoch *et al*., 1991). To date, a large number of C-to-U RNA editing sites have been reported and collected into REDIdb 3.0 and PED database (Lo Giudice *et al*., 2018; Li *et al*., 2019). Most of C-to-U RNA editing events occurred in the protein-coding regions to change the corresponding amino acid sequences (Yan *et al*., 2018). Occasionally, some C-to-U RNA editing sites were located in the non-coding regions, which affected RNA alternative splicing and protein translation efficiency (Castandet *et al*., 2010). Collectively, increasing evidences indicated that C-to-U RNA editing were involved in various biological processes, such as chloroplast biogenesis in *Arabidopsis* (Yu *et al*., 2009), retrograde signaling in *Arabidopsis* (Zhao *et al*., 2019), heat tolerance in *Arabidopsis* (Huang *et al*., 2022), and normal fruit development in tomato (Li *et al*., 2023).

Currently, several software tools are developed to identify chloroplast RNA editing sites. For example, PREP-Cp first translated a given RNA sequence into a protein sequence, and then made predictions based on homologous sequences alignment (Mower, 2009). Nevertheless, PREP-Cp could not deal with non-coding RNA sequence and synonymous cytidines for codon degeneracy (Edera *et al*., 2021). Another software, according to sequence conservation, CURE-Chloroplast identified the potential C-to-U RNA editing sites for seed plants via a multiple sequences alignment (Du *et al*., 2009). This shortcoming is that the flanking sequences surrounding the C-to-U RNA editing sites were not always conserved but some species-specific sites still exist (Okuda and Shikanai, 2012; Shikanai, 2015). Furthermore, the growing studies revealed that RNA editing have tissue/condition-specific patterns, which was studied by RNA-seq data (Kim *et al*., 2016; Zhang *et al*., 2020). At the same time, some RNA editing pipelines were built to detect RNA editing sites from RNA-seq data. Although considerable efforts were made, accurate prediction of RNA editing sites were usually affected by many factors, such as the quality of input RNA-seq data and genome mapping strategies (Diroma *et al*., 2019). Especially, for non-model plants, if no reference plastid genomes are available, thus, we are unable to perform RNA-seq analysis for identification of RNA editing sites.

In this study, we developed PlantC2U by utilizing a convolutional neural network (CNN) for genome sequence-dependent identification of plastid C-to-U RNA editing sites in plants. Also, PlantC2U could further check modification status of C-to-U RNA editing sites identified by RNA-seq analysis. Based on CNN, PlantC2U outperformed conventional random forest (RF) and support vector machine (SVM) model. Interestingly, we observed a similar sequence context surrounding the RNA editing sites to that in *Arabidopsis* (Chu and Wei, 2020*b*), suggesting that editing factors possibly operates in different species editing systems (Oldenkott *et al*., 2020). Beside*s*, our results showed that the preference of overall C-to-U RNA editing level across the different tissues in mangrove plant *Kandelia obovata*. Additional possible functionality of PlantC2U is that PlantC2U can systematically estimate the key regions for RNA-guided RNA editing. In summary, we developed a PlantC2U online tool and presented RNAeditDB, available online at https://jasonxu.shinyapps.io/RNAeditDB/, the first genome-wide identification of plastid C-to-U RNA editing in mangrove species.

## Materials and methods

### Datasets

To build PlantC2U for identification of RNA editing in the plastid of plants, we obtained chloroplast RNA editing events from REDIdb 3.0 (http://srv00.recas.ba.infn.it/redidb/index.html) (Lo Giudice *et al*., 2018), which included cDNA position of RNA editing modification, the corresponding genomic sequence and cDNA sequence. Then, we extracted base information of RNA editing site from genomic sequence and cDNA sequence based on position of RNA editing modification, respectively, and kept only genomic sequence with C-to-T substitution (C-to-U in mRNA; C-to-T in DNA) for sequence feature extraction. As for mix-bases, for instance M (Adenine/Cytidine), symbols for mix-bases from genomic sequence were replaced with Ns symbols. In general, the flanking sequences of the base modification are sufficient to recognize RNA editing events (Shikanai, 2015; Tac *et al*., 2021). Therefore, for each RNA editing site, we extracted the flanking sequence of different lengths nucleotides (nt) from a cytidine.

In brief, PlantC2U program required the input sequences to keep a fixed length (default: 100 nt). To meet the demand of PlantC2U, we filled the both ends of those shorter sequences with Ns to match the fixed length. After that, the cross-species sequences without any redundancy were used as positive instances. At the same time, we also extracted the corresponding sequences on upstream and downstream region of cytidines without RNA editing modification in the edited genes but excluded in the positive dataset, which were used to create our primary negative samples. Each input sequence consisted of 100 nucleotides centered on each cytidine. We adopted one-hot encoding method to convert each nucleotide from 100-nt sequence into a binary vector, such as A (1, 0, 0, 0, 0), C (0, 1, 0, 0, 0), G (0, 0, 1, 0, 0), T (0, 0, 0, 1, 0), N (0, 0, 0, 0, 1). Thus, each 100-nt sequence was represented by a 5×100 binary matrix with rows corresponding to A, C, G, T and N as the input to PlantC2U.

### Selection of negative samples

In order to improve our prediction of RNA editing, we evaluated the effects of selecting negative samples on model. Firstly, we used a similar strategy described in previous study to cluster nucleotide sequences from positive samples and primary negative samples (Macosko *et al*., 2015; James *et al*., 2018). Here, each positive sample and primary negative sample sequence was represented by a matrix of *k*-mer count that consisted of different short subsequences of length *k* (Kirk *et al*., 2018). That is, a matrix of *k*-mer count, in which rows are *k*-mers, and columns are nucleotide sequences indices, was used to perform data normalization, data scaling, dimensional reduction, and nucleotide sequences clustering using the Seurat v.4.1.1 R package (Hao *et al*., 2021*a*). We used K-Nearest Neighbors (KNN) and Louvain algorithm from FindNeighbors and FindClusters function to perform nucleotide sequences clustering based on the first 10 principal components.

Secondly, we chose different datasets as negative instances using two different sampling methods: (i) Random Negative Sequences (RNS1) from primary negative samples; (ii) Random Negative Sequences (RNS2) from different sequences clusters. Importantly, in order to evaluate the effect of the ratio of positive and negative instances, the ratio of the positive and negative instances was set as 1:1, 1:2, 1:3, 1:4 and 1:5, respectively. Finally, we merged the positive and negative instances for machine learning. The performance of models for selection of negative instances were then compared.

### Model architecture

The deep learning model, PlantC2U, was constructed based on the Keras v.2.9.0 R package (https://tensorflow.rstudio.com/) with the TensorFlow v.2.9.0 library as a backend in R v4.2.1. We utilized a similar CNN architecture used in the previous study (Wang and Wang, 2019), which included 2 one-dimensional convolution layer (Conv1D), a max pooling layer, and a flatten layer below:

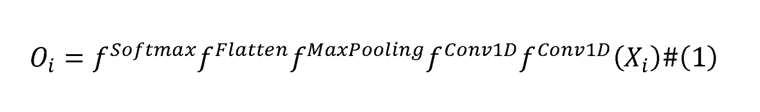

The x_i_ represented a 5×100 binary matrix, which was fed into the first Conv1D layer with 5 channels (A, C, G, T and N), 12 kernel size, 1 stride, rectified liner unit (Relu) activation function, and 0% dropout. The second Conv1D layer with 5 channels, 6 kernel size, 1 stride, Relu activation function was used to learn high dimensional features. Then, the max pooling layer with 5 pool size, and 4 strides was used to reduce the dimensionality of the output from the second Conv1D layer with 50% dropout. The output from the max pooling layer with 50% dropout was flattened into one vector, which fully connected to the last layer. The output of the last layer with two nodes (o_i_) was to calculate the prediction probability for identification of RNA editing using the nonlinear softmax activation function.

### Model performance evaluation

For positive and negative instances, we randomly divided them into two subsets: 75% for training and 25% for testing. The deep learning model was trained on the training dataset using a 10-fold cross-validation method, and then the best model was chosen to evaluate the performance based on the testing dataset. The following summary statistics metrics (Du *et al*., 2009) of Sensitivity, Specificity, Positive Predictive Value (PPV), Accuracy, Balanced Accuracy (BA) and the Matthew’s Correlation Coefficient (MCC), were used to evaluate the performance of PlantC2U:

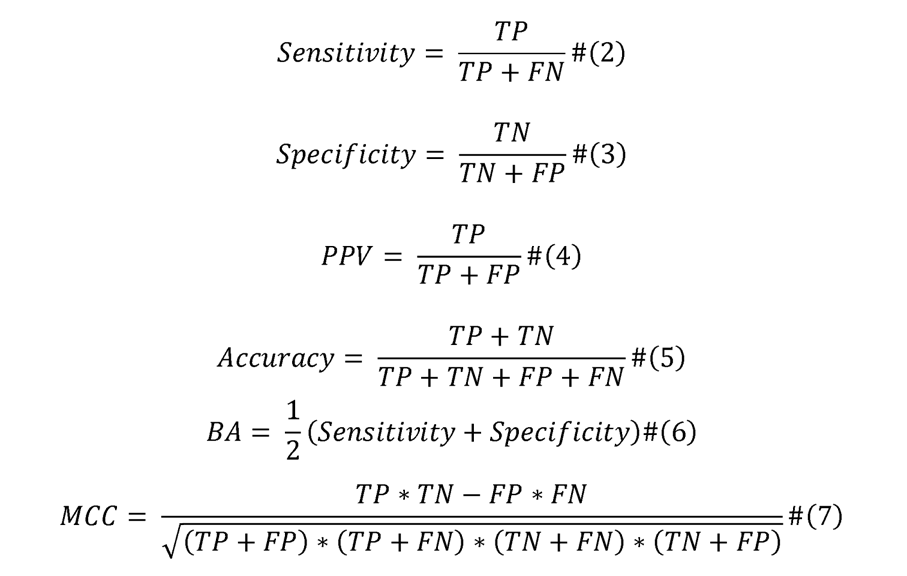

As mentioned above, TP, FP, TN and FN represented the number of true positives, false positives, true negatives and false negatives, respectively. In addition, comparison of PlantC2U with several machine learning methods for classification were tested, including random forest (RF), support vector machine (SVM). The area under the receiver operating characteristic curve (AUC-ROC) and precision-recall curve (AUC-PRC) were also used to evaluate the performance of models.

### Application of PlantC2U to mangrove plant *K. obovata* RNA-seq datasets

We downloaded the RNA-seq raw data in eight tissues of *K. obovata* from NCBI SRA database (NCBI Resource Coordinators, 2018) with accession SRP123536 for identification of RNA editing sites, including flower, fruit, leaf, pistil, root, sepal, stamen, and stem tissue (Su *et al*., 2019). Then, the raw reads were extracted and cleaned using SRA toolkit (Version: 2.11.2; https://github.com/ncbi/sra-tools) and Trim Galore software (Version: 0.6.6; https://github.com/FelixKrueger/TrimGalore), respectively. The high-quality reads in FASTQ format were mapped to the plastid genome of *K. obovata* using HISAT2 software with strand-specific mode based on RNA-seq library construction strategy (Kim *et al*., 2019). After the alignments, the SAM files were converted, sorted and deduplicated using SAMtools (v1.9) (Danecek *et al*., 2021) and sambamba (v0.8.1) (Tarasov *et al*., 2015), respectively.

For identification of RNA editing sites, the BAM files were used to perform SNV calling using REDItools (Picardi and Pesole, 2013; Flati *et al*., 2020). To obtain the reliable RNA editing sites we found, we retained only SNVs with at least 5% editing level and 10 covered read supports, and removed SNVs with multiple variant types. Subsequently, the PlantC2U can be optionally applied to distinguish C-to-U editing sites from possible SNPs called by RNA-seq analysis.

### Construction of the mangrove RNAeditDB

To build a comprehensive database of plastid C-to-U RNA editing sites in mangrove species, we also obtained the raw RNA-seq datasets in other mangrove species from the National Genomics Data Center (NGDC) under accession number CRA004363 using Edge Turbo Client (https://ngdc.cncb.ac.cn/ettrans/?filePath=/gsa/CRA004363). The corresponding plastid genomes were downloaded from the Chloroplast Genome Information Resource (CGIR; https://ngdc.cncb.ac.cn/cgir) and NCBI (Hua *et al*., 2022). As mentioned above, we adopted a standardized workflow to uniformly process the raw RNA-seq datasets.

The expression levels of plastid genes were quantified by featureCounts and StringTie2 (Liao *et al*., 2014; Kovaka *et al*., 2019, page 2). The DEseq2 R package was used to perform gene counts normalization, and then detect differential expression genes (DEGs) (Love *et al*., 2014, page 2). The DEGs were determined by *P* value < 0.05 and the log-scale fold change (|log_2_FC| >= 0.5). And we used Wilcoxon test to identify differential RNA editing sites with statistical significance (*P* value < 0.1) between control condition and test condition. Of note, due to different RNA-seq library construction strategies, strand-specific or non-strand-specific mode was implemented in the process of the reads mapping and SNVs calling. Besides, we used the SnpEff and SnpSift program to annotate the RNA editing sites (Cingolani *et al*., 2012; Cingolani, 2022).

## Results

### Construction of PlantC2U

As demonstrated in **Figure 1**, PlantC2U was developed for identification of C-to-U RNA editing sites using CNN. In the REDIdb 3.0, C-to-U editing sites were the most common RNA editing type (**Table S1**). We collected a total of 1,961 C-to-U RNA editing sites of 640 RNA edited genes from 88 organisms (**Table S2**). In this study, we used RNA editing sites six times more than previous studies (319 and 301) as positive samples for model construction (Du *et al*., 2009; Mower, 2009). Compared to positive samples, 102,324 cytidines without RNA editing in the same edited genes were extracted for negative samples (**Table S2**). We focused on using only nucleotides sequences that flanking cytidines to predict whether cytidines can be edited to uridines.

**Figure 1.**
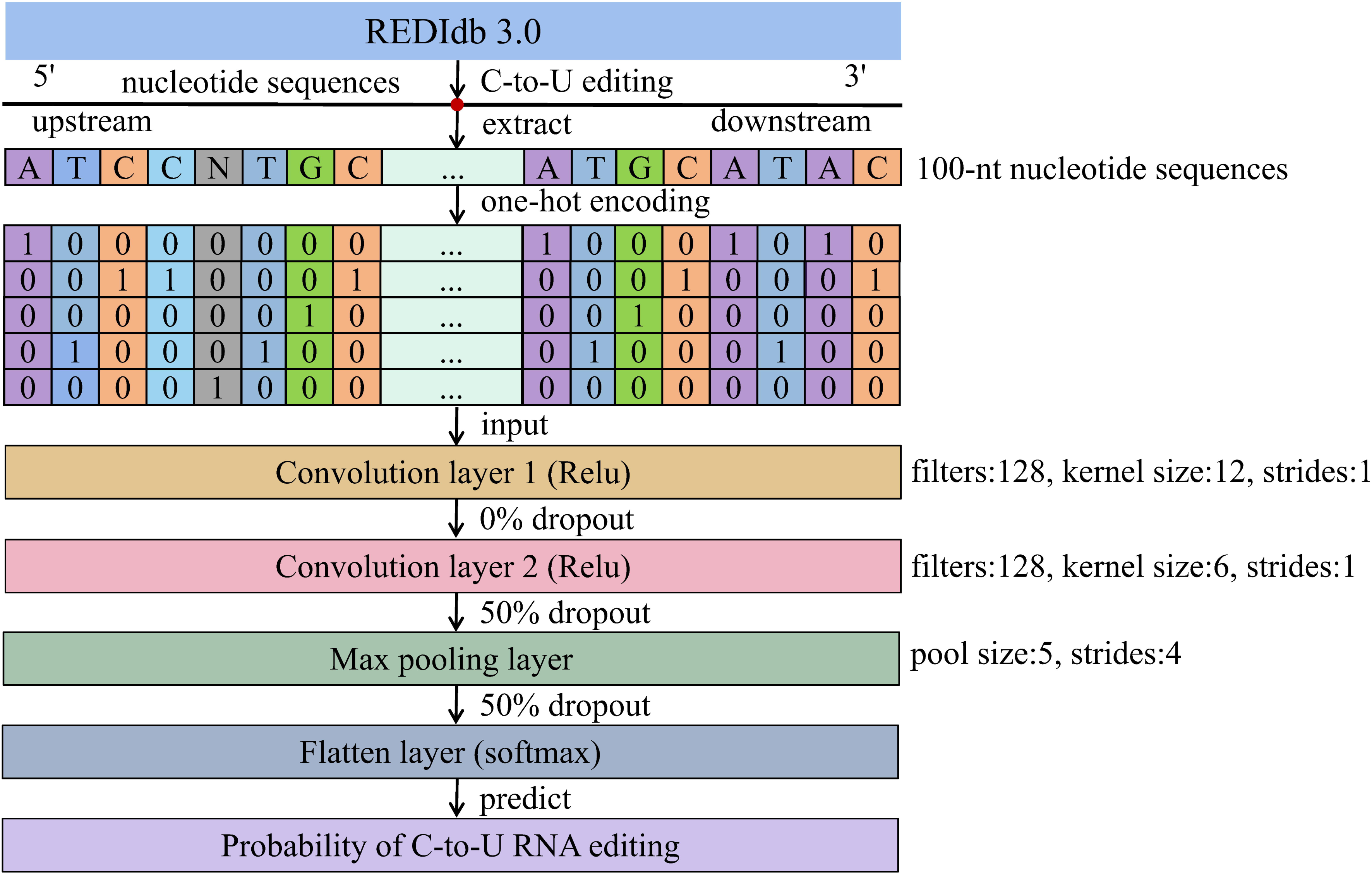
A schematic diagram of prediction of plastid C-to-U RNA editing sites. The 100-nt flanking sequences surrounding the target cytidines are extracted and converted into a 5×100 binary matrix as the input for PlantC2U. The CNN architecture includes 2 one-dimensional convolution layers (Conv1D), a max pooling layer, and a flatten layer. The nonlinear softmax activation function is used to calculate the prediction probability of C-to-U RNA editing.

Subsequently, we have tested several combinations of the flanking sequence of different length and the different ratios of positive and negative instances to determine the best training model using a 10-fold cross-validation strategy. In other words, our pipeline of PlantC2U consisted of three steps: (1) Extraction of the flanking sequence; (2) Evaluation of different negative samples (3) Comparison with other machine learning approaches.

### Choice of negative samples affects model performance

More and more studies have showed that the choice of negative samples can effectively improve the performance of machine learning models (Cheng *et al*., 2017, 2018; Song *et al*., 2020). We extracted the flanking sequence from cytidines between 20-nt and 100-nt sequence in steps length of 10-nt sequence to perform PlantC2U model. For selected random negative samples (RNS1) from primary negative samples, to some extent, we found that the lower ratio of positive and negative instances, the better behaviors of machine learning models (**Figure 2A**). Therefore, we provided the MCC and PPV values to evaluate the performance of models for unbalanced datasets. **Figure 2A** and **Figure S1A** suggested that the 60-nt or 70-nt flanking sequence made the model better based on MCC and PPV. For 70-nt flanking sequence, the corresponding mean PPV and MCC value of model were 0.88 and 0.60, respectively (**Table S3**).

**Figure 2.**
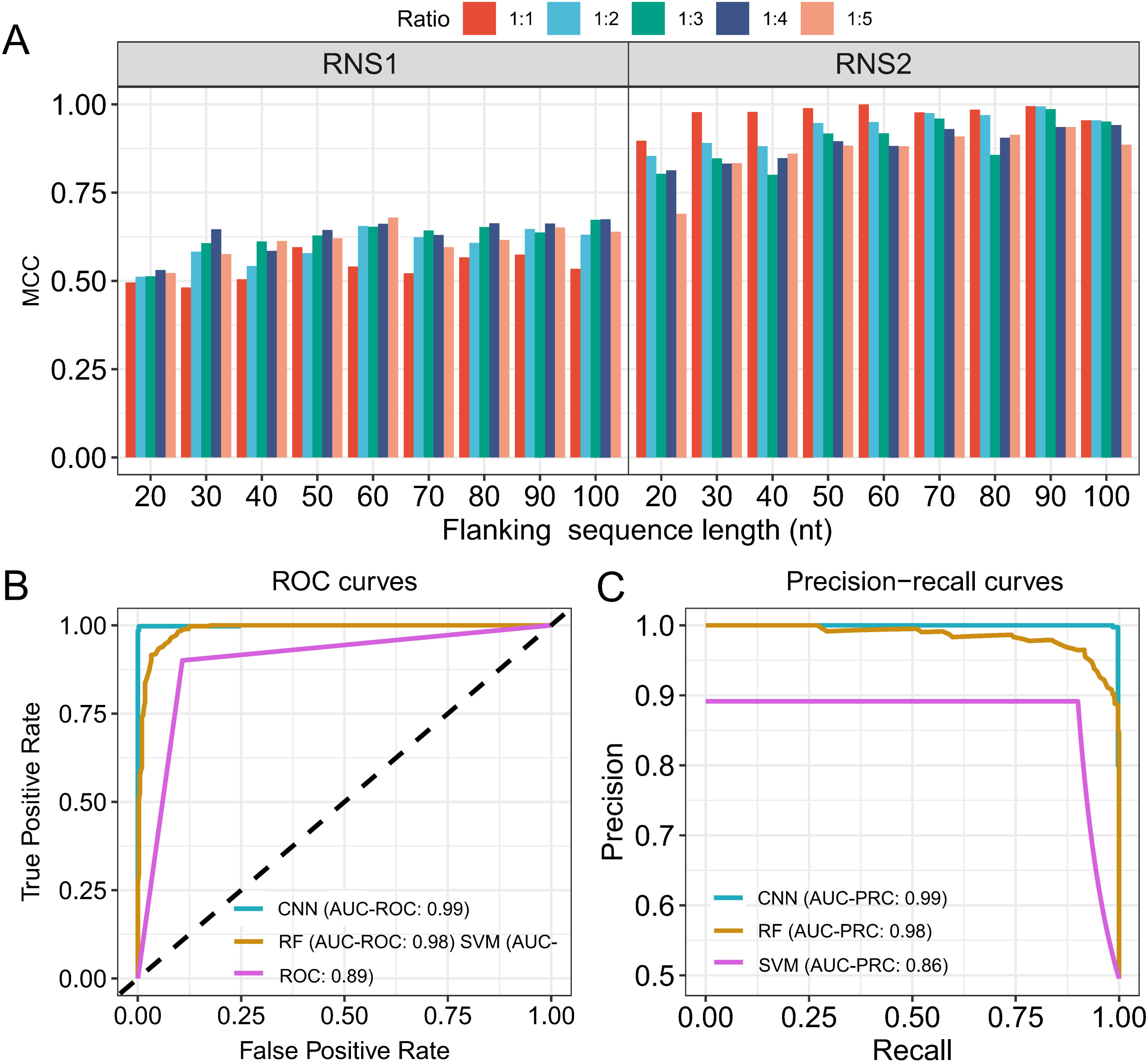
The evaluation of model performance under different combinations of negative and positive samples. (A) The model performance based on the Matthew’s Correlation Coefficient (MCC) using RNS1 and RNS2 as negative samples, respectively. RNS1 represents random negative sequences from primary negative samples; RNS2 represents random negative sequences from different sequences clusters; The Ratio is positive-negative samples ratio. (B) The receiver operating characteristic (ROC) curves show the better performance of convolutional neural network (CNN) than the random forest (RF) and support vector machine (SVM) models. (C) Evaluation of model performance based on the area under the precision-recall curves (AUC-PRC).

However, for selected random negative samples (RNS2) from different sequences clusters, we observed a opposite phenomenon that the higher ratio of positive and negative instances, the better behaviors of machine learning models (**Figure 2A; Figure S1B**). Therefore, we also provided the balanced accuracy (BA) values to evaluate the performance of models for balanced datasets. We found that the 90-nt flanking sequence in combination with the 1:1 ratio of positive and negative instances made the model better based on BA values (**Figure S1B**). Overall, the PlantC2U model with the 90-nt flanking sequence achieved more than 95% sensitivity and 99% specificity, and the corresponding mean BA and MCC value of model were 0.98 and 0.97, respectively (**Table S4**). These results indicated that utilization of RNS2 can be a robust and effective approach to obtain better model (**Figure S2A**). As a result, we chose the 90-nt flanking sequence and the 1:1 ratio of positive and negative instances for the following analysis.

### Comparison of machine learning approaches and sequence context analysis

In this work, we also tested random forest (RF) and support vector machine (SVM) learning models, and compared model prediction ability of RNA editing modification among them. As expected, three models (RF, SVM and CNN) performed more superior than random guess (AUC-ROC: 0.5, diagonal line). Of which, CNN model had the highest AUC-ROC at 0.99 and the highest AUC-PRC at 0.99 based on the testing data (**Figure 2B and Figure 2C**). In summary, PlantC2U outperformed RF and SVM machine learning models in this study. Our results suggested that the flanking sequences surrounding the target cytidines were sufficient to recognize whether the cytidines will be edited into uridines.

In addition, more and more studies demonstrated that C-to-U editing sites preferentially occurred in a sequence context manner (Di Giorgio *et al*., 2020; Chu and Wei, 2020*b*). Therefore, in order to identify enriched motif, we extracted the 5-nt sequences flanking the C-to-U editing sites to perform sequence context analysis. We then calculated the frequency of A, C, G and T in the upstream and downstream region flanking the C-to-U editing sites. Compared to randomly selected unedited cytidines sites, there was significantly different sequence context. And the sequence logo showed that the flanking sequences of C-to-U editing sites mainly enriched in the ‘TGTTTCAATTG’ motif, in contrast, the flanking sequences of unedited cytidines sites mainly enriched in the ‘TGGTTCTTTTT’ motif (**Figure S2B and Figure S2C**). These sequence context revelated that a potential mechanism to recognize C-to-U editing by *cis*-regulatory elements (Okuda and Shikanai, 2012; Lesch *et al*., 2022). A recent study reported that three or more mutations near the editing sites would lead to an abolishment of RNA editing event (Liu *et al*., 2021). To our knowledge, the PlantC2U program can be applied to assess the effect of mutations on C-to-U editing status by learning *cis*-regulatory features.

### Characterization of C-to-U RNA editing landscape in *K. obovata*

Another utility of PlantC2U is to filter out possible SNPs from C-to-U editing sites called by RNA-seq analysis. In the present study, we detected 5,863 RNA editing sites from 24 RNA-seq datasets of eight tissues in the plastids of *K. obovata* (**Figure 3**), including 1,468 C-to-U RNA editing sites (C-to-U in mRNA; C-to-T in DNA), while 6.47% (95/1,468) C-to-U RNA editing sites possibly resulted from SNPs (**Table S5**). We focused on C-to-U RNA editing type, which was the most common RNA editing type in *K. obovata* (**Figure S3**). This result was consistent with previous observations in other plants, such as *Arabidopsis thaliana* and rice (Zhang *et al*., 2017; Lo Giudice *et al*., 2018; Chu and Wei, 2020*b*). By our large analysis, we observed that C-to-U editing levels differently distributed across eight tissues (**Figure 4A and Figure S4**). Most of C-to-U editing sites were located in the protein-coding regions. Among them, 766 editing sites were nonsynonymous substitutions, and even 160 editing sites generated a new stop codon (**Table S6**). These results indicated that plastid RNA editing modifications resulted in change of amino acid sequences.

**Figure 3.**
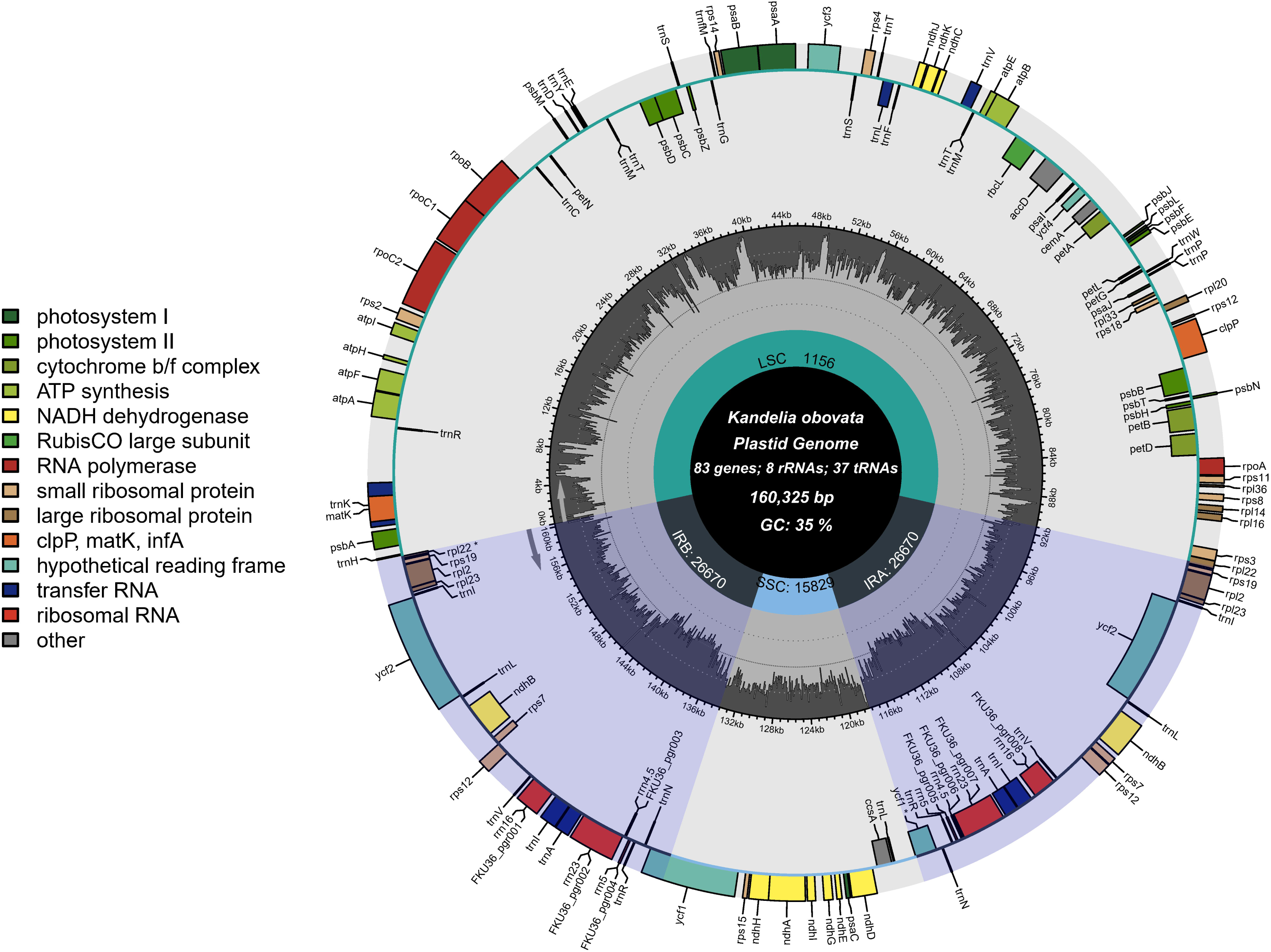
Representative map of the plastid genome of mangrove plant *K. obovata*, including 83 genes, 8 rRNAs, 37 tRNAs, and 35% GC content. IRA and IRB indicate inverted repeat regions; LSC is large single copy, SSC is small single copy.

**Figure 4.**
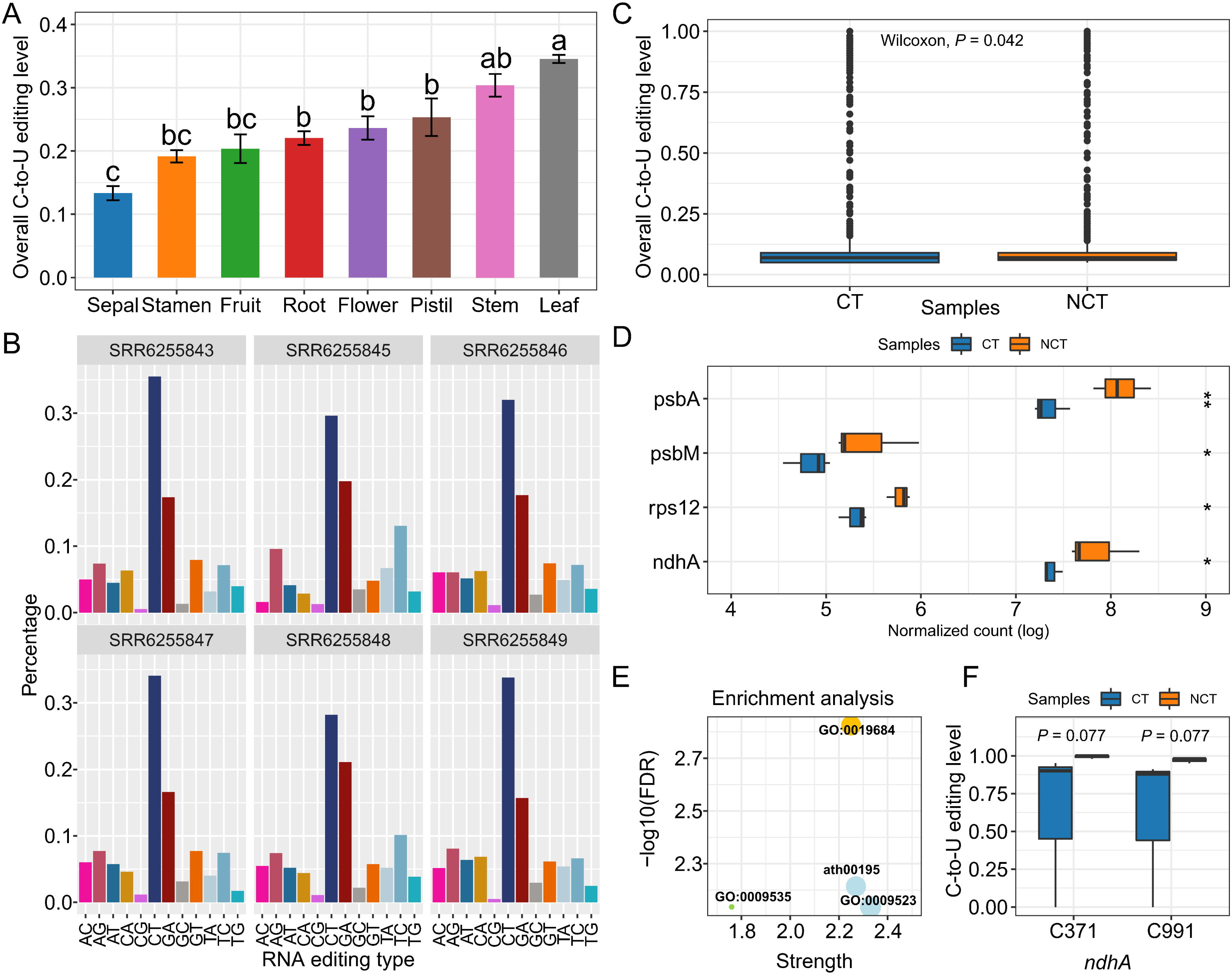
Identification of plastid RNA editing sites in *K. obovata*. (A) Overall C-to-U RNA editing level in different *K. obovata* tissues. Values indicate mean ±SD of three biological replicates. The significant difference of RNA editing level were represented by different lowercases through one-way ANOVA (*P* < 0.05). (B) SNV identified from *K. obovata* leaf transcriptomes. The bar plots show the percentage of each SNV type (e.g., C>T, CT) identified in each *K. obovata* transcriptome. (C) The box plots show the overall C-to-U RNA editing level in cold tolerant (CT) and non-cold tolerant (NCT) transcriptomes. (D) The box plots of the expression level of four genes (*psbA*, *psbM*, *rps12* and *ndhA*) in CT and NCT transcriptomes. ’*’ represents 0.01 < *P* ≤ 0.05, ’**’ represents 0.001 < *P* ≤ 0.01. (E) The functional enrichment analysis of four genes (*psbA*, *psbM*, *rps12* and *ndhA*) using STRING website tool. (F) The C-to-U RNA editing level of C371 and C991 sites from *ndhA* in CT and NCT transcriptomes.

To further investigate C-to-U RNA editing events, we identified 2,263 plastid RNA editing sites from 6 RNA-seq datasets in *K. obovata* leaf tissues (3 cold tolerant (CT) samples and 3 non-cold tolerant (NCT) samples). We also used PlantC2U program to detect 45 possible SNPs from 731 C-to-U RNA editing sites (**Table S7**). Admittedly, the C-to-U RNA editing type was the most common RNA editing type (**Figure 4B**). Moreover, the global C-to-U editing level of cold tolerant samples was significantly decreased when compared to the non-cold tolerant samples (*P* value < 0.05; **Figure 4C**). Further functional enrichment analysis (Szklarczyk *et al*., 2019) (https://cn.string-db.org/) showed that four DEGs (*psbA*, *psbM*, *rps12* and *ndhA*) were significantly enriched in “Photosynthesis, light reaction”, Photosynthesis, “Photosystem II”, and Chloroplast thylakoid membrane (**Figure 4D and Figure 4E; Table S8**).

Next, we focused on C-to-U RNA editing sites on these four DEGs, namely *psbA*, *psbM*, *rps12* and *ndhA*, and found two pivotal RNA editing sites (C371 and C991) on *ndhA*. The editing level of C371 and C991 sites had a significant difference between CT and NCT (*P* value < 0.1; **Figure 4F**). Further investigation demonstrated that C371 would create a new stop codon and C991 caused a nonsynonymous change of amino acid from Proline (Pro) to Threonine (Thr) for *ndhA*. Furthermore, these two RNA editing sites were less likely to be caused by SNPs through PlantC2U checking. Our results implied that the C-to-U RNA editing events may play an important role in the chilling stress tolerance of *K. obovata*.

### Identification of C-to-U RNA editing sites in other 24 mangrove species

In this study, we provided the first genome-wide characterization of plastid C-to-U RNA editing events in mangrove species. Here, a total of 50 RNA-seq datasets across four tissues (flower, fruit, leaf and root) of 24 mangrove species were obtained for identification of C-to-U RNA editing (**Table S9**). In general, we uniformly processed files from the raw data to BAM files, and then performed *de novo* identification of C-to-U RNA editing, at last, annotated the RNA editing sites (**Figure 5A**). After alignment, the mean reference-guided alignment rate was about 2.98% (**Figure S5A**). As a result, we identified 5,229 RNA editing sites from the gene body of plastid genes, including 2,381 C-to-U RNA editing sites, which were the most abundant type of RNA editing (**Figure S5B**). In addition, the mean number of C-to-U RNA editing sites for each sample was approximate 50 (**Figure 5B**). The majority of C-to-U RNA editing sites (∼90%) were located in the protein-coding region, including 1,760 nonsynonymous codon changes and 147 sites resulted in new stop codons (**Table S10**). Of which, we found 74 possible SNPs by PlantC2U program. Taken together, all C-to-U RNA editing sites we identified were stored in the RNAeditDB.

**Figure 5.**
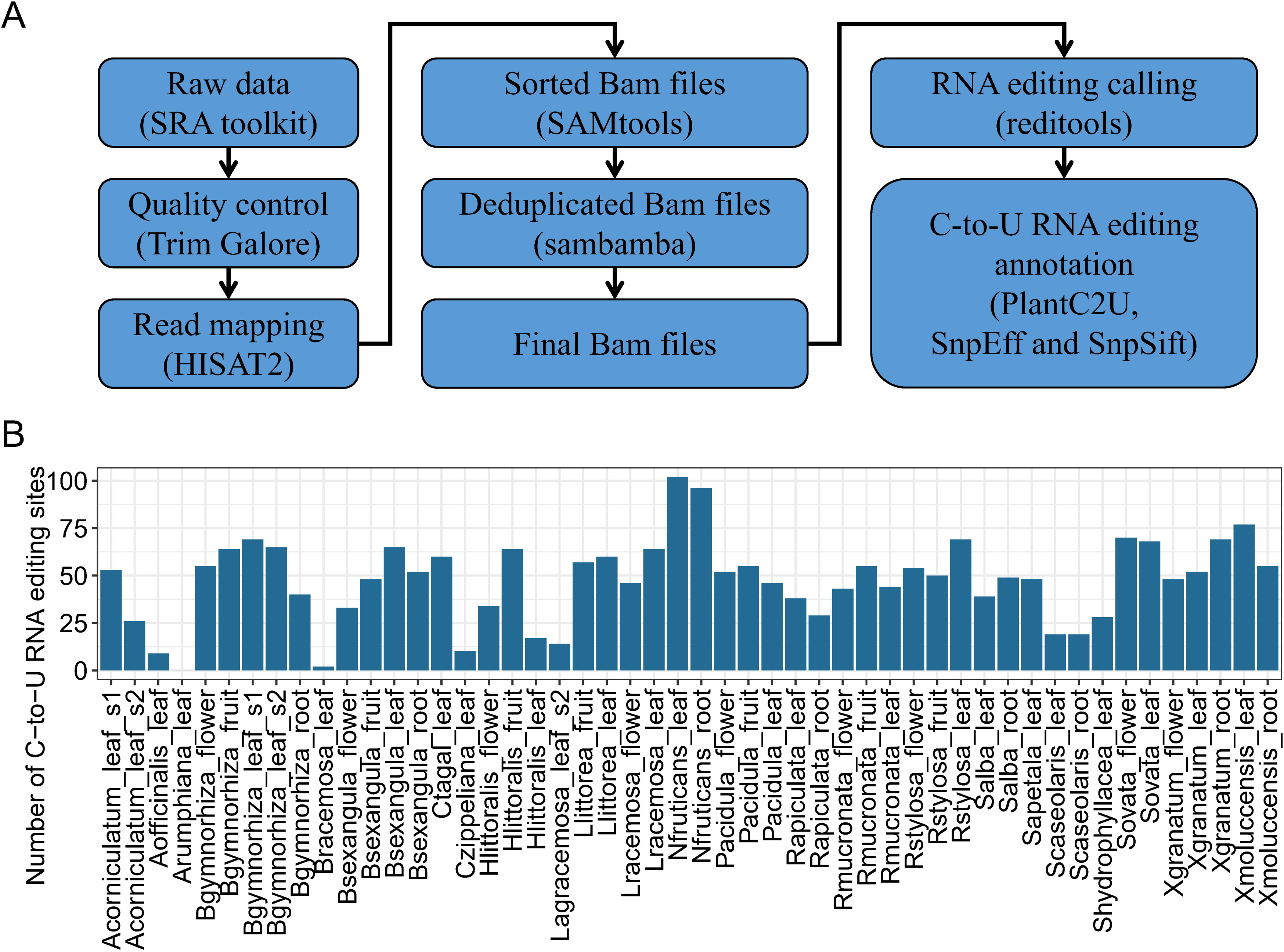
A workflow for identification and annotation of plastid RNA editing sites. (A) A pipeline includes quality control of raw sequencing data, clean read mapping, RNA editing sites calling, and C-to-U RNA editing sites annotation. (B) Number of C-to-U RNA editing sites identified from different tissues of 24 mangrove species.

### Construction of PlantC2U and RNAeditDB

We developed an PlantC2U online tool via shinyApp (https://www.shinyapps.io/), which was used to predict plastid C-to-U RNA editing sites in plant species (https://jasonxu.shinyapps.io/RNAeditDB/). Users could paste the nucleotide sequences in FASTA format, and select flanking sequence length and ratio of positive and negative samples to obtain the probability of a potential C-to-U RNA editing site (**Figure 6**). In addition, we built RNAeditDB to store all C-to-U RNA editing sites in mangrove species.

**Figure 6.**
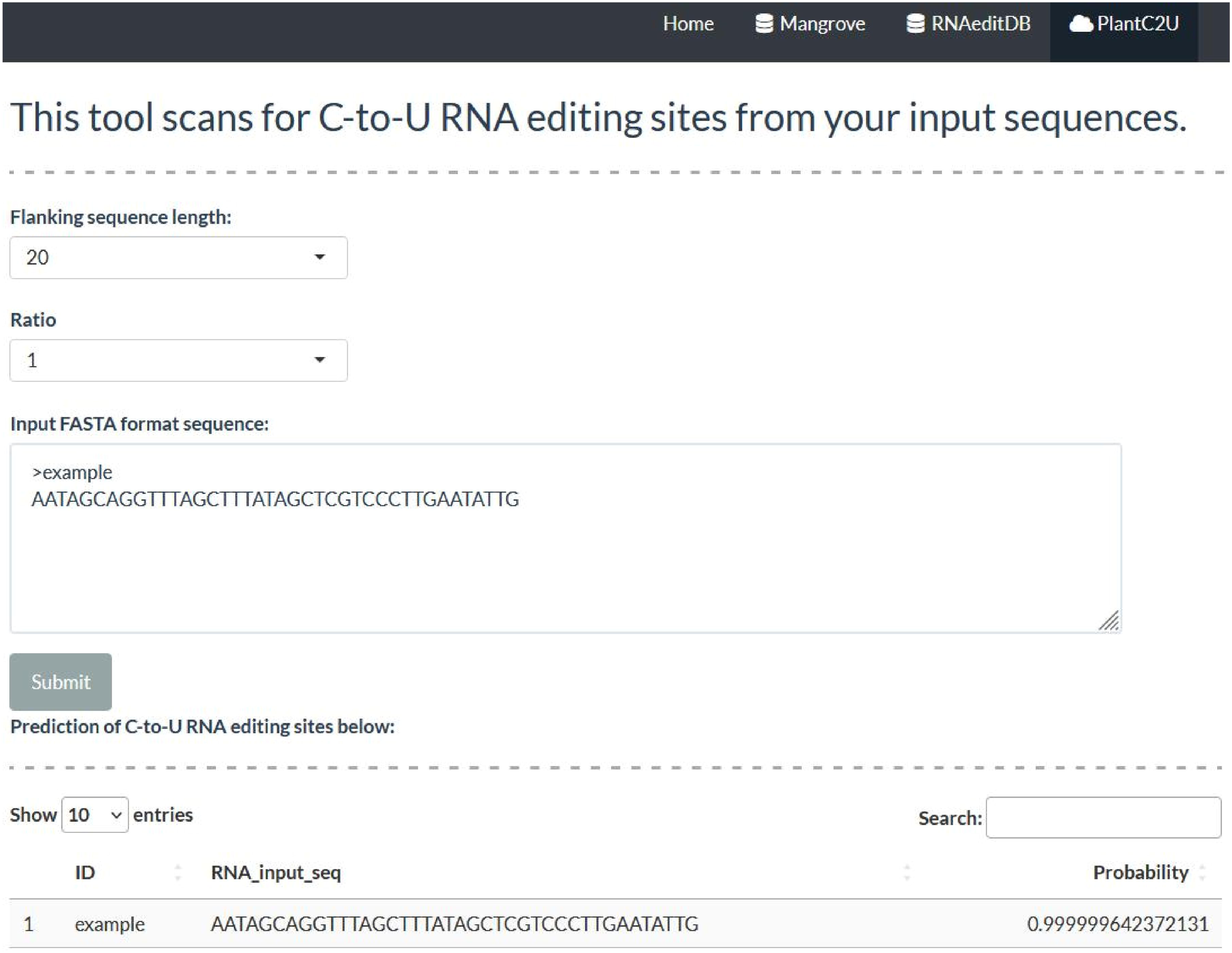
Interface of the online predictor, PlantC2U, on plants plastid C-to-U RNA editing sites.

## Discussion

RNA editing by C-to-U determination is a prominent post-transcriptional modification process that modifies the genetic code in the RNA level, which is commonly found in the plastids and mitochondria in plants (Small *et al*., 2020; Hao *et al*., 2021*b*). More and more studies elucidated the important roles of RNA editing events in the plant growth and development (He *et al*., 2018; Zhang *et al*., 2020). In particular, the rapid development of RNA deep sequencing technologies provided unprecedented opportunities to detect RNA editing events from the massive transcriptome data. However, for most software packages, the main limitation of RNA editing sites prediction is that there was no matched DNA-seq data or available known SNPs dataset for non-model species (Zhang and Xiao, 2015; Sun *et al*., 2016; Wang *et al*., 2016; Flati *et al*., 2020). Although REDItools and RED software could explore RNA editing events using RNA-seq alone, there were still false positives and false negatives (Sun *et al*., 2016). Therefore, in the present study, we developed a sequence-based PlantC2U program to further filter C-to-U RNA editing sites identified by RNA-seq analysis, partly due to variants such as SNPs.

Compared to other studies (Du *et al*., 2009; Mower, 2009), we obtained more C-to-U RNA editing sites in plants for model training, including 1,961 positive instances and 102,324 negative instances. Nevertheless, our dataset was still severely imbalanced. Here, we tested two types of negative samples: (i) RNS1, which were randomly selected from 102,324 negative instances; and (ii) RNS2, which were negative instances from 71 clusters after clustering all positive and negative instances (**Table S2**). We demonstrated that the average model performance using RNS2 as negative samples was better than using RNS1. One possible reason was that sequences clustering method could find some very similar negative instances as positive instances, which had low frequency of RNA editing (Yan *et al*., 2018). Our results revelated that negative samples were randomly selected from the same sequence cluster with a relatively low proportion of positive instances would effectively improve model performance (Cheng *et al*., 2017).

Importantly, we also included a flanking sequence length and positive-to-negative instances ratio selection step, where we found that 90-nt flanking sequence and the 1:1 ratio had a superior prediction ability of C-to-U RNA editing. Based on the 90-nt flanking sequence of C-to-U RNA editing sites, we found a similar sequence context to that in *Arabidopsis thaliana* (Chu and Wei, 2020*b*), even in viral (Di Giorgio *et al*., 2020). Lesch *et al*. have confirmed that plant mitochondrial RNA editing factors could perform targeted C-to-U editing of nuclear transcripts in human cells (Lesch *et al*., 2022). Thus, it is of great interest whether plant plastid RNA editing factors could play similar roles in human nuclear transcripts, due to sequence context. In addition, more evidences have supported that point mutation near the editing sites would change RNA editing level (Freyer *et al*., 1997; Sapiro *et al*., 2015; Sharma and Baysal, 2017). In our model, using computationally different mutation combinations near the editing sites, we may analyze the effects of each mutation combination on RNA editing level, thereby identifying potential *cis*-regulatory elements.

In this study, we systematically identified 2,381 C-to-U RNA editing sites in the plastid of 24 mangrove species. The circular plastid genomes of mangrove species were 148-168kb in size, indicating a ∼0.06% editing frequency in the plastid genome, which was higher than a previous study (Sasaki *et al*., 2001). It is largely because mangroves plants grow in extreme intertidal environments with high salinity, hypoxia and other abiotic stresses (Nizam *et al*., 2022). And RNA editing events were usually beneficial to plant developmental process, including adaption to environment changes (Yan *et al*., 2018). Interestingly, consistent with previous study, we found that reducing C-to-U RNA editing rates may contribute to the cold adaption in mangrove plant *K. obovata* (Chu and Wei, 2020*a*; Zhang *et al*., 2020). However, it’s necessary to validate experimentally our findings and that would be conducted in our following study.

It is also noteworthy that there are some limitations of PlantC2U program. First, the PlantC2U could distinguish possible SNPs from C-to-U RNA editing sites based on corresponding flanking sequences and RNA-seq data, but this inference needs to be validated by cDNA sequence analysis. Second, *trans*-factors for RNA editing and RNA secondary structure would affect C-to-U RNA editing (Shikanai, 2015; Shi *et al*., 2016), thereby these characteristics should be considered for model training in the future. Finally, the PlantC2U program was constructed based on the plastid RNA editing sites, and whether we can predict C-to-U RNA editing events in the mitochondria of plants by PlantC2U. Taken together, we developed an PlantC2U online tool, and built the RNAeditDB web interface to predict and show the plastid C-to-U RNA editing sites.

## Supplementary data

**Figure S1.** The evaluation of model performance under different combinations of negative and positive samples.

**Figure S2.** Sequences clustering and nucleotide context flanking the C-to-U editing sites.

**Figure S3**. SNV identified from 24 RNA-seq datasets of eight tissues in the plastid of *K. obovata*. The bar plots show the percentage of each SNV type (e.g., C>T, CT).

**Figure S4.** The RNA editing level of plastid genes across eight tissues of *K. obovata*.

**Figure S5.** SNV identified in 50 mangrove transcriptomes.

**Table S1.** The detailed information of plastid RNA editing sites collected in REDIdb 3.0.

**Table S2.** The summary of the plastid C-to-U RNA editing sites dataset.

**Table S3.** The performance of PlantC2U in independent test based on the different flanking sequence and the ratio of positive and negative instances from primary negative samples.

**Table S4.** The performance of PlantC2U in independent test based on the different flanking sequence and the ratio of positive and negative instances from sequence clusters.

**Table S5.** Identification of plastid C-to-U editing sites across eight tissues in *K. obovata*.

**Table S6.** Annotation and functional prediction of plastid C-to-U editing sites on the protein-coding genes in *K. obovata*.

**Table S7.** Identification of plastid C-to-U editing sites in *K. obovata* leaf tissues under chilling events.

**Table S8.** The functional enrichment analysis of four DEGs (*psbA*, *psbM*, *rps12* and *ndhA*)

**Table S9.** The detailed information of 50 RNA-seq datasets, covering 4 tissues from 24 mangrove species.

**Table S10.** Identification of plastid C-to-U editing sites in other 24 mangrove species.

### Abbreviations

C-to-U: cytidine to uridine transition
CNN: convolutional neural network
RF: random forest
SVM: support vector machine
KNN: K-Nearest Neighbors
Conv1D: one-dimensional convolution layer
Relu: rectified liner unit
PPV: Positive Predictive Value
BA: Balanced Accuracy
MCC: Matthew’s Correlation Coefficient
CT: cold tolerant samples
NCT: non-cold tolerant samples
AUC-ROC: the area under the receiver operating characteristic curve
AUC-PRC: the area under the precision-recall curve

## Acknowledgments

We appreciate anonymous reviewers and the editor for the insightful comments and valuable suggestions.

## Author contributions

The project was conceived and directed by H-LZ. Data analysis and construction of RNAeditDB website were performed by CX and JL. L-YS contributed to discussions on machine learning models. JL and Z-JG produced tables and figures. JL collected data. S-WS and L-DZ checked images clarity. The manuscript was written by CX, and revised by H-LZ. All authors discussed interpretations and commented on the manuscript. All authors have read and approved the final manuscript.

## Data availability

All data supporting the findings of this study are available in Figshare (https://doi.org/10.6084/m9.figshare.22802885.v1).

## Reference

1. Brennicke A, Marchfelder A, Binder S. 1999. RNA editing. FEMS Microbiology Reviews 23, 297–316.

2. Castandet B, Choury D, Bégu D, Jordana X, Araya A. 2010. Intron RNA editing is essential for splicing in plant mitochondria. Nucleic Acids Research 38, 7112–7121.

3. Cheng Z, Huang K, Wang Y, Liu H, Guan J, Zhou S. 2017. Selecting high-quality negative samples for effectively predicting protein-RNA interactions. BMC Systems Biology 11, 9.

4. Cheng Z, Zhou S, Wang Y, Liu H, Guan J, Chen Y-PP. 2018. Effectively identifying compound-protein interactions by learning from positive and unlabeled examples. IEEE/ACM Transactions on Computational Biology and Bioinformatics 15, 1832–1843.

5. Chu D, Wei L. 2020a. Reduced C-to-U RNA editing rates might play a regulatory role in stress response of Arabidopsis. Journal of Plant Physiology 244, 153081.

6. Chu D, Wei L. 2020b. Systematic analysis reveals cis and trans determinants affecting C-to-U RNA editing in Arabidopsis thaliana. BMC Genetics 21, 98.

7. Cingolani P. 2022. Variant Annotation and Functional Prediction: SnpEff. Methods in Molecular Biology (Clifton, N.J.) 2493, 289–314.

8. Cingolani P, Patel VM, Coon M, Nguyen T, Land SJ, Ruden DM, Lu X. 2012. Using Drosophila melanogaster as a model for genotoxic chemical mutational studies with a new program, SnpSift. Frontiers in Genetics 3, 35.

9. Danecek P, Bonfield JK, Liddle J, et al. 2021. Twelve years of SAMtools and BCFtools. GigaScience 10, giab008.

10. Di Giorgio S, Martignano F, Torcia MG, Mattiuz G, Conticello SG. 2020. Evidence for host-dependent RNA editing in the transcriptome of SARS-CoV-2. Science Advances 6, eabb5813.

11. Diroma MA, Ciaccia L, Pesole G, Picardi E. 2019. Elucidating the editome: Bioinformatics approaches for RNA editing detection. Briefings in Bioinformatics 20, 436–447.

12. Du P, Jia L, Li Y. 2009. CURE-Chloroplast: A chloroplast C-to-U RNA editing predictor for seed plants. BMC Bioinformatics 10, 135.

13. Edera AA, Small I, Milone DH, Sanchez-Puerta MV. 2021. Deepred-Mt: Deep representation learning for predicting C-to-U RNA editing in plant mitochondria. Computers in Biology and Medicine 136, 104682.

14. Flati T, Gioiosa S, Spallanzani N, Tagliaferri I, Diroma MA, Pesole G, Chillemi G, Picardi E, Castrignanò T. 2020. HPC-REDItools: A novel HPC-aware tool for improved large scale RNA-editing analysis. BMC Bioinformatics 21, 353.

15. Freyer R, Kiefer-Meyer M-C, Kössel H. 1997. Occurrence of plastid RNA editing in all major lineages of land_plants. Proceedings of the National Academy of Sciences of the United States of America 94, 6285–6290.

16. Hao Y, Hao S, Andersen-Nissen E, et al. 2021a. Integrated analysis of multimodal single-cell data. Cell 184, 3573–3587.e29.

17. Hao W, Liu G, Wang W, et al. 2021b. RNA editing and its roles in plant organelles. Frontiers in Genetics 12.

18. He P, Xiao G, Liu H, Zhang L, Zhao L, Tang M, Huang S, An Y, Yu J. 2018. Two pivotal RNA editing sites in the mitochondrial atp1mRNA are required for ATP synthase to produce sufficient ATP for cotton fiber cell elongation. The New Phytologist 218, 167–182.

19. Hiesel R, Wissinger B, Schuster W, Brennicke A. 1989. RNA editing in plant mitochondria. Science (New York, N.Y.) 246, 1632–1634.

20. Hoch B, Maier RM, Appel K, Igloi GL, Kössel H. 1991. Editing of a chloroplast mRNA by creation of an initiation codon. Nature 353, 178–180.

21. Hua Z, Tian D, Jiang C, Song S, Chen Z, Zhao Y, Jin Y, Huang L, Zhang Z, Yuan Y. 2022. Towards comprehensive integration and curation of chloroplast genomes. Plant Biotechnology Journal 20, 2239–2241.

22. Huang C, Liu D, Li Z-A, Molloy DP, Luo Z-F, Su Y, Li H-O, Liu Q, Wang R-Z, Xiao L-T. 2022. The PPR protein RARE1-mediated editing of chloroplast accD transcripts is required for fatty acid biosynthesis and heat tolerance in Arabidopsis. Plant Communications 4, 100461.

23. James BT, Luczak BB, Girgis HZ. 2018. MeShClust: An intelligent tool for clustering DNA sequences. Nucleic Acids Research 46, e83.

24. Kim M, Hur B, Kim S. 2016. RDDpred: a condition-specific RNA-editing prediction model from RNA-seq data. BMC genomics 17 Suppl 1, 5.

25. Kim D, Paggi JM, Park C, Bennett C, Salzberg SL. 2019. Graph-based genome alignment and genotyping with HISAT2 and HISAT-genotype. Nature Biotechnology 37, 907–915.

26. Kirk JM, Kim SO, Inoue K, et al. 2018. Functional classification of long non-coding RNAs by k-mer content. Nature Genetics 50, 1474–1482.

27. Kovaka S, Zimin AV, Pertea GM, Razaghi R, Salzberg SL, Pertea M. 2019. Transcriptome assembly from long-read RNA-seq alignments with StringTie2. Genome Biology 20, 278.

28. Lesch E, Schilling MT, Brenner S, Yang Y, Gruss OJ, Knoop V, Schallenberg-Rüdinger M. 2022. Plant mitochondrial RNA editing factors can perform targeted C-to-U editing of nuclear transcripts in human cells. Nucleic Acids Research 50, 9966–9983.

29. Li J, Wang K, Yang Y, et al. 2023. SlRIP1b is a global organellar RNA editing factor, required for normal fruit development in tomato plants. The New Phytologist 237, 1188–1203.

30. Li M, Xia L, Zhang Y, et al. 2019. Plant editosome database: A curated database of RNA editosome in plants. Nucleic Acids Research 47, D170–D174.

31. Liao Y, Smyth GK, Shi W. 2014. featureCounts: An efficient general purpose program for assigning sequence reads to genomic features. Bioinformatics 30, 923–930.

32. Liu X, Sun T, Shcherbina A, Li Q, Jarmoskaite I, Kappel K, Ramaswami G, Das R, Kundaje A, Li JB. 2021. Learning cis-regulatory principles of ADAR-based RNA editing from CRISPR-mediated mutagenesis. Nature Communications 12, 2165.

33. Lo Giudice C, Pesole G, Picardi E. 2018. REDIdb 3.0: A Comprehensive Collection of RNA Editing Events in Plant Organellar Genomes. Frontiers in Plant Science 9, 482.

34. Love MI, Huber W, Anders S. 2014. Moderated estimation of fold change and dispersion for RNA-seq data with DESeq2. Genome Biology 15, 550.

35. Macosko EZ, Basu A, Satija R, et al. 2015. Highly parallel genome-wide expression profiling of individual cells using nanoliter droplets. Cell 161, 1202–1214.

36. Mower JP. 2009. The PREP suite: Predictive RNA editors for plant mitochondrial genes, chloroplast genes and user-defined alignments. Nucleic Acids Research 37, W253–259.

37. NCBI Resource Coordinators. 2018. Database resources of the National Center for Biotechnology Information. Nucleic Acids Research 46, D8–D13.

38. Nizam A, Meera SP, Kumar A. 2022. Genetic and molecular mechanisms underlying mangrove adaptations to intertidal environments. iScience 25.

39. Okuda K, Shikanai T. 2012. A pentatricopeptide repeat protein acts as a site-specificity factor at multiple RNA editing sites with unrelated cis-acting elements in plastids. Nucleic Acids Research 40, 5052–5064.

40. Oldenkott B, Burger M, Hein A-C, Jörg A, Senkler J, Braun H-P, Knoop V, Takenaka M, Schallenberg-Rüdinger M. 2020. One C-to-U RNA editing site and two independently evolved editing factors: Testing reciprocal complementation with DYW-Type PPR proteins from the moss Physcomitrium (Physcomitrella) patens and the flowering plants Macadamia integrifolia and Arabidopsis. The Plant Cell 32, 2997–3018.

41. Picardi E, Pesole G. 2013. REDItools: High-throughput RNA editing detection made easy. Bioinformatics 29, 1813–1814.

42. Sapiro AL, Deng P, Zhang R, Li JB. 2015. Cis regulatory effects on A-to-I RNA editing in related Drosophila species. Cell reports 11, 697–703.

43. Sasaki Y, Kozaki A, Ohmori A, Iguchi H, Nagano Y. 2001. Chloroplast RNA editing required for functional acetyl-CoA carboxylase in plants. The Journal of Biological Chemistry 276, 3937–3940.

44. Sharma S, Baysal BE. 2017. Stem-loop structure preference for site-specific RNA editing by APOBEC3A and APOBEC3G. PeerJ 5, e4136.

45. Shi X, Germain A, Hanson MR, Bentolila S. 2016. RNA recognition motif-containing protein ORRM4 broadly affects mitochondrial RNA editing and impacts plant development and flowering. Plant Physiology 170, 294–309.

46. Shikanai T. 2015. RNA editing in plants: Machinery and flexibility of site recognition. Biochimica et Biophysica Acta (BBA) - Bioenergetics 1847, 779–785.

47. Small ID, Schallenberg-Rüdinger M, Takenaka M, Mireau H, Ostersetzer-Biran O. 2020. Plant organellar RNA editing: What 30 years of research has revealed. The Plant Journal: For Cell and Molecular Biology 101, 1040–1056.

48. Song Q, Lee J, Akter S, Rogers M, Grene R, Li S. 2020. Prediction of condition-specific regulatory genes using machine learning. Nucleic Acids Research 48, e62.

49. Su W, Ye C, Zhang Y, Hao S, Li QQ. 2019. Identification of putative key genes for coastal environments and cold adaptation in mangrove Kandelia obovata through transcriptome analysis. The Science of the Total Environment 681, 191–201.

50. Sun Y, Li X, Wu D, Pan Q, Ji Y, Ren H, Ding K. 2016. RED: A Java-MySQL software for identifying and visualizing RNA editing sites using rule-based and statistical filters. PLOS ONE 11, e0150465.

51. Szklarczyk D, Gable AL, Lyon D, et al. 2019. STRING v11: Protein-protein association networks with increased coverage, supporting functional discovery in genome-wide experimental datasets. Nucleic Acids Research 47, D607–D613.

52. Tac HA, Koroglu M, Sezerman U. 2021. RDDSVM: Accurate prediction of A-to-I RNA editing sites from sequence using support vector machines. Functional & Integrative Genomics 21, 633–643.

53. Tarasov A, Vilella AJ, Cuppen E, Nijman IJ, Prins P. 2015. Sambamba: Fast processing of NGS alignment formats. Bioinformatics 31, 2032–2034.

54. Wang Z, Lian J, Li Q, Zhang P, Zhou Y, Zhan X, Zhang G. 2016. RES-Scanner: a software package for genome-wide identification of RNA-editing sites. GigaScience 5, 37.

55. Wang J, Wang L. 2019. Deep learning of the back-splicing code for circular RNA formation. Bioinformatics 35, 5235–5242.

56. Yan J, Zhang Q, Yin P. 2018. RNA editing machinery in plant organelles. Science China Life Sciences 61, 162–169.

57. Yu Q-B, Jiang Y, Chong K, Yang Z-N. 2009. AtECB2, a pentatricopeptide repeat protein, is required for chloroplast transcript accD RNA editing and early chloroplast biogenesis in Arabidopsis thaliana. The Plant Journal: For Cell and Molecular Biology 59, 1011–1023.

58. Zhang Z, Cui X, Wang Y, Wu J, Gu X, Lu T. 2017. The RNA Editing Factor WSP1 Is Essential for Chloroplast Development in Rice. Molecular Plant 10, 86–98.

59. Zhang A, Jiang X, Zhang F, Wang T, Zhang X. 2020. Dynamic response of RNA editing to temperature in grape by RNA deep sequencing. Functional & Integrative Genomics 20, 421–432.

60. Zhang Q, Xiao X. 2015. Genome sequence-independent identification of RNA editing sites. Nature Methods 12, 347–350.

61. Zhao X, Huang J, Chory J. 2019. GUN1 interacts with MORF2 to regulate plastid RNA editing during retrograde signaling. Proceedings of the National Academy of Sciences 116, 10162–10167.

